# Profiling of linear B-cell epitopes against human coronaviruses in pooled sera sampled early in the COVID-19 pandemic

**DOI:** 10.1101/2024.02.29.582263

**Authors:** Emil Bach, Mustafa Ghanizada, Nikolaj Kirkby, Søren Buus, Thomas Østerbye

## Abstract

**Background:** Antibodies play a key role in the immune defence against infectious pathogens. Understanding the underlying process of B cell recognition is not only of fundamental interest; it supports important applications within diagnostics and therapeutics. Whereas the nature of conformational B cell epitope recognition is inherently complicated, linear B cell epitopes offer a straightforward approach that potentially can be reduced to one of peptide recognition.

**Methods:** Using an overlapping peptide approach representing the entire proteomes of the seven main coronaviruses known to infect humans, we analysed sera pooled from eight PCR-confirmed COVID-19 convalescents and eight pre-pandemic controls. Using a high-density peptide microarray platform, 13-mer peptides overlapping by 11 amino acids were in situ synthesised and incubated with the pooled primary serum samples, followed by development with secondary fluorochrome-labelled anti-IgG and -IgA antibodies. Interactions were detected by fluorescence detection. Strong Ig interactions encompassing consecutive peptides were considered to represent “high-fidelity regions” (HFRs). These were mapped to the coronavirus proteomes using a 60% homology threshold for clustering.

**Results:** We identified 333 human coronavirus derived HFRs. Among these, 98 (29%) mapped to SARS-CoV-2, 144 (44%) mapped to one or more of the four circulating common cold coronaviruses (CCC), and 54 (16%) cross-mapped to both SARS-CoV-2 and CCCs. The remaining 37 (11%) mapped to either SARS-CoV or MERS-CoV. Notably, the COVID-19 serum was skewed towards recognising SARS-CoV-2-mapped HFRs, whereas the pre-pandemic was skewed towards recognising CCC-mapped HFRs. In terms of absolute numbers of linear B cell epitopes, the primary targets are the ORF1ab protein (60%), the spike protein (21%), and the nucleoprotein (15%) in that order; however, in terms of epitope density the order would be reversed.

**Conclusion:** We identified linear B cell epitopes across coronaviruses, highlighting pan-, alpha-, beta-, or SARS-CoV-2-corona-specific B cell recognition patterns. These findings could be pivotal in deciphering past and current exposures to epidemic and endemic coronavirus. Moreover, our results suggest that pre-pandemic anti-CCC antibodies may cross-react against SARS-CoV-2, which could explain the highly variable outcome of COVID-19. Finally, the methodology used here offers a rapid and comprehensive approach to high-resolution linear B-cell epitope mapping, which could be vital for future studies of emerging infectious diseases.

## Introduction

As of November 2023, the coronavirus disease 2019 (COVID-19) pandemic has resulted in more than 676 million infections and 6.8 million deaths worldwide (COVID-19 Map - Johns Hopkins Coronavirus Resource Center); a mortality rate of 1% [1]. The causative agent of COVID-19 and its genome was determined in January 2020, identifying a novel severe acute respiratory syndrome coronavirus 2 (SARS-CoV-2) [2]. SARS-CoV-2 belongs to the Coronaviridae family, wherein seven virus strains infect humans, all causing respiratory tract infections [3]. These seven human coronaviruses (HCoVs) encompass two genera: Five are in the Betacoronavirus: HCoV-OC43, HCoV-HKU1 (sub-strains N1, N2, and N5), SARS-CoV, SARS-CoV-2, and Middle East respiratory syndrome coronavirus (MERS-CoV). Two are in the Alphacoronavirus genus: HCoV-NL63 and HCoV-229E [4]. OC43, HKU1, NL63, and 229E, are collectively known as the common cold coronavirus (CCC). The CCCs are endemic and responsible for 5-30% of common cold cases [3]. They cause mild respiratory symptoms and are considered non-fatal. In contrast, the epidemic SARS-CoV (emerged in 2002) and MERS-CoV (emerged in 2012) exhibited high mortality rates of approximately 9% and 36%, respectively.

Coronaviruses are enveloped, positive-sense RNA viruses. The SARS-CoV-2 genome contains 15 open reading frames (ORFs) encoding (1) the structural proteins making up the envelope of the virion, the spike (S), envelope (E), membrane (M), and nucleocapsid (N) proteins; and (2) the accessory proteins found in the polyproteins ORF1a and ORF1ab together with ORFs 3a, 3b, 6, 7a, 7b, 8, 9b, 9c, and 10 [5]. This genomic architecture is similar among the HCoVs, which all share the polyproteins (ORF1a/ab) and structural proteins (S, E, M, and N) [5]. At the sequence level, their nucleotide sequence identities range from 47-80% [6]. Comparing S proteins, SARS-CoV-2 has the highest similarity with SARS-CoV, with a nucleotide sequence identity of 80.4% and protein sequence identity of 77.3% [6]. The HCoV S proteins facilitate their entrance into their host cells via interactions with host cell surface proteins [3,5,7], making them ideal targets for vaccines aimed at inducing neutralising antibodies. Indeed, the S protein is the main target in many SARS-CoV-2 vaccines in use and under development [8,9]. It follows that mutations in the S protein are particularly problematic for the longevity of the immunity induced by these vaccines.

Since its emergence, SARS-CoV-2 has accumulated genomic changes such as mutations, indels (insertions and deletions), and recombinations, resulting in novel variants [10]. Most proteomic mutations have occurred in the ORF1ab protein, followed by the S protein and then the N protein [11–13]. The SARS-CoV-2 Omicron ’complex’ (PANGO lineages BA.1, BA.2, BA.3, BA.4, BA.5) developed more than 15 mutations in the S protein receptor-binding domain (RBD) and multiple genomic changes in the amino-terminal domain (NTD) [10]. These mutations have led to antigenic changes in the S protein, allowing variants in the Omicron complex to escape neutralising antibodies in sera from first-generation vaccinees and pre-Omicron convalescents. The dominant variant for the first half of 2023 was the XBB and its sub-lineages, originating from a recombination between two BA.2 sub-lineages [10,14]. The XBB variant exhibited even higher antibody evasion than its parental strains [15]. However, the XBB variant is currently in the process of being outcompeted by EG.5, a new descendant lineage of XBB.19.2 [14]. This new variant shows even greater resistance than XBB to neutralisation by sera from vaccinees with or without breakthrough infections with other circulating subvariants compared to the original sub-variants in the Omicron complex [16–19]. Thus, the evasion of humoral immunity appears to be an important driver of SARS-CoV-2 evolution [15].

Antibodies are produced by B cells and the specific regions of an antigen that they recognise are known as B cell epitopes. The identification of virus-specific B cell epitopes is important for rational approaches to prevent, treat, diagnose, and understand virus infections. In general, antibodies against protein antigens can be broadly categorized into those that are specific for conformational (discontinuous) and those that are specific for linear (continuous) B cell epitopes; conformational B cell epitopes are composed of amino acids that are brought into proximity through the folded tertiary structure of a protein antigen, whereas linear B cell epitopes consist of a sequential string of amino acids from the primary structure of a protein antigen.

Vital for their infectivity, viruses use ligand(s) to find and attach to specific target(s) on appropriate host cells (e.g., SARS-CoV-2 uses its spike protein to specifically attach to ACE2 on the target cell). Antibodies that bind to a virus ligand and interfere with the infective virus-host cell interaction are likely to have virus-neutralising activity. For many natural infections, the corresponding neutralising B cell epitope is thought to be conformational. This is highlighted in the recent design of SARS-CoV-2 vaccines. Here, the vaccines have predominantly been targeted against the infective pre-fusion structure of the virus’s spike protein. To enhance the vaccine’s efficacy, specific mutations have been introduced to stabilise this crucial structure.

Whether a neutralising antibody recognises a conformational or linear B cell epitope has important theoretical and practical considerations. About 90% of B cell epitopes are conformational [20]. Unfortunately, identifying conformational B cell epitopes experimentally tends to involve demanding and low throughput structural studies such as X-ray crystallography of antibody-antigen complexes, NMR or deuterium exchange in the presence or absence of antibody, directed mutagenesis, etc. (reviewed in [21]). Machine learning-based predictors of conformational B cell epitopes have been introduced [22,23] and are continuously improving [24–26]. Representing conformational epitopes (e.g. for formulation in vaccines or diagnostics) typically requires recombinant expression in organisms that allow proper folding and post-translational modifications. About 10% of B cell epitopes are linear [20], which typically can be represented by oligopeptides.

Comparatively, it is straightforward to identify linear B cell epitopes since large peptide libraries can be generated synthetically (e.g. as peptide microarrays) or recombinantly (e.g. as phage display libraries). Using overlapping peptide strategies, it is possible to cover entire pathogen proteomes, and using single substitution analysis, it is possible to delineate the length of individual B cell epitopes and define the underlying motifs with several discrete positions where certain amino acids are required, allowed, or disallowed.

We have pioneered a photolithographic peptide microarray technology, which can produce over a hundred thousand peptides per microarray [27]. Using this platform to screen for linear B cell epitopes, we have observed that the typical length of a linear B cell epitope is 5-10 amino acids (range 4-12 amino acids) and that it typically features a compound specificity with a few very dominant positions allowing very few amino acid substitutions, some less dominant position allowing several amino acids and a few “spacer” positions allowing all amino acids [27–29]. Notably, our studies revealed that the specificity of linear B cell epitope recognition demonstrated remarkable conformational sensitivity, enabling precise discrimination among closely related amino acids, such as serine versus threonine, glutamine versus asparagine etc. [27]. Here, we have used this high-density peptide microarray platform to examine and compare the linear B cell epitope repertoire against human coronavirus in pooled sera from eight COVID-19 convalescents sampled around March 2020 (i.e. early in the COVID-19 pandemic) and eight pre-pandemic individuals sampled before October 2019. We report early pandemic and pre-pandemic IgG and IgA responses against linear B cell epitopes from SARS-CoV-2, SARS-CoV, and MERS-CoV, as well as the four human common cold coronaviruses.

## Methods

### Early COVID-19 pandemic patient and healthy pre-pandemic control serum pools

Pools of anonymised serum from early pandemic, COVID-19-positive convalescent donors or pre-pandemically sampled negative control donors were obtained in March 2020 from the University Hospital of Copenhagen. The COVID-19-positive status of the COVID-19 convalescent donors was confirmed using a commercial IVD-certified RT-PCR test to detect the presence of SARS-CoV-2 RNA in oropharyngeal swaps. Serum from eight COVID-19 convalescents and eight pre-pandemic donors were selected randomly from a larger panel of potential donors, de-identified, and pooled, thus creating anonymous pools of COVID-19 positive and negative donors.

### High-density peptide microarrays

As a non-overlapping source of negative control peptides (NP), we randomly generated 3900 random 13 amino acid peptides using the amino acid frequencies of the investigated proteomes. As an overlapping source of negative control peptides (EP), we extracted the proteome of Zaire Ebola virus (strain Mayinga-76, EBOZM), which has not circulated in Denmark. Thus, we did not expect any Danish human serum pool to contain antibodies against EBOZM-derived peptides. As a source of positive control peptides (PP), we extracted the proteome of human cytomegalovirus (HCMV, strain AD169). Due to the strong reactivity and high seroprevalence of HCMV in the human population [30], we expected (and later confirmed) that any human serum pool would contain antibodies against HCMV-derived peptides.

To design experimental libraries of human coronavirus-derived peptides (CP), we extracted the proteomes of the seven human coronaviruses (HCoVs) from the UniProt Knowledgebase [31]: the common cold coronaviruses (CCC), including the alpha coronaviruses 229E and NL63 and the beta coronaviruses OC43 and HKU1(N1 isolate), the epidemic HCoVs MERS-CoV and SARS-CoV, and the novel coronavirus, SARS-CoV-2. All the virus proteomes were represented by overlapping peptide libraries consisting of 13 amino acid long peptides (Table 1) tiled by 2 amino acids (i.e., overlapping by 11 amino acids) and filtered for sequence redundancy. A high-density peptide microarray design was generated, distributing the CPs, EPs and PPs in triplicate and the NPs in duplicate randomly across 12 virtual sectors using proprietary software (PepArray, Schafer-N). Peptides were synthesised by Schafer-N (Copenhagen) on amino-functionalized glass microscope slides using a maskless photolithographic synthesis strategy as described elsewhere [27]. After de-protection and washing, the peptide microarrays were used to detect anti-peptide-specific antibodies.

**Table 1.** Number of peptides from each organism on the peptide microarrays.

| Organism (Abbreviation) | Peptide Count* |
| --- | --- |
| Zaire ebolavirus (strain Mayinga-76) (EBOZM) | 2417 |
| Human cytomegalovirus (strain AD169) (HCMV) | 31388 |
| Severe acute respiratory syndrome coronavirus 2 (SARS-CoV-2) | 6330 |
| Human SARS coronavirus (SARS-CoV) | 6353 |
| Middle East respiratory syndrome-related coronavirus (isolate United Kingdom/H123990006/2012) (MERS-CoV) | 4933 |
| Human coronavirus HKU1 (isolate N1) (HCoV-HKU1(N1)) | 4998 |
| Human coronavirus OC43 (HCoV-OC43) | 5164 |
| Human coronavirus NL63 (HCoV-NL63) | 4689 |
| Human coronavirus 229E (HCoV-229E) | 4605 |
| Total* | 66581 |

\* Total is the union of all peptide sets derived from the viral proteomes.

### Detecting the peptide-specific binding of IgG and IgA antibodies

Peptide microarrays were washed in PBS supplemented with 0.1% BSA, 0.1% Triton X-100 and incubated with serum pools from convalescent COVID-19 patients or pre-pandemic donors diluted 1:100 in PBS (0.1% BSA, 0.1% Triton X-100) for 2 hours at room temp, washed in PBS (0.1% BSA, 0.1% Triton X-100), and developed by detecting bound IgG or IgA with appropriate fluorochrome-labelled secondary antibodies: 1 µg/ml goat anti-human IgG-Cy3 (Sigma, Cat# C2571) or goat anti-human IgA-Cy5 (Jackson ImmunoResearch, Cat# 109-175-011), respectively. Following post-incubation washes, the microarrays were spin-dried and read in a microarray laser scanner (INNOSCAN 900, Innopsys, France). Finally, the proprietary PepArray software was used to quantify antibody-binding to each peptide at 8-bit resolution.

### Identifying peptides binding antibodies with high-fidelity

Staining artefacts were automatically identified by the proprietary software, PepArray (Schafer-N, Copenhagen, Denmark), and the corresponding peptide signals were removed from the final dataset. The remaining data from the high-density peptide microarray were analysed using the R statistical language version 4.3.0 [32]. For each serum pool and each Ig isotype, the arithmetic mean signal for each virus-derived peptide and NP was calculated, and the background was subtracted using the median signal of all 3900 random peptides (NPs) to account for non-specific binding (any background corrected value ≤ 0 were recoded to 0, and a value of 1 was added to all signals). The empirical cumulative distribution (Equation 1) of the NP background-corrected signals was used to assess the probability (p-value) that antibody-bindings signals recorded for virus-derived peptides could be fortuitous (i.e., non-specific).

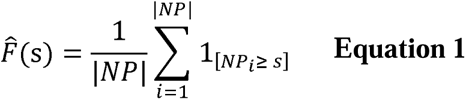

Where NP indicates the antibody-binding signals of peptides in the NP set, and s indicates an antibody-binding signal of any virus-derived peptide. The virus-derived peptide sequences were mapped to their parent proteins. Each virus-derived peptide sequence was pooled with its neighbouring sequences overlapping with at least 11 positions in the parent protein using the R package, IRanges version 2.36.0 [33]. Antibody-binding fidelity for each virus-derived peptide was assessed by combining its p-value with the p-values of its pooled neighbours using Fisher’s combined probability test (Equation 2).

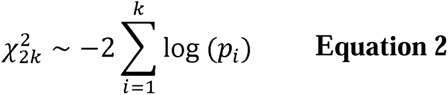

Where k indicates the number of pooled peptides and pi the p-value of peptide i in pool k derived from Equation 1. We reasoned that our overlapping peptide library strategy involving 13-mer peptides overlapping by 11 amino acids would ensure that linear B cell epitopes, largely being less than 10 amino acids long [27], would appear in at least two consecutive overlapping peptides. Moreover, to focus our search on B cell epitopes that yielded signals that were clearly above the background, we imposed a minimum signal requirement of 20% of the maximum signal obtained. Thus, we set a threshold for high-fidelity antibody binding for each virus peptide pair at a Fisher’s combined p-value ≤ 0.001 and a standardised and cleaned signal value ≥ 20% of the maximum standardised and background-corrected signal value measured for the serum pool and Ig-type in question. This effectively means that any solitary peptide with a high binding signal will be considered a false positive and assigned to the non-binder group which consists of all remaining virus-derived peptides outside the high-fidelity group. The number of peptides determined to have high-fidelity antibody-binding (HFPs), are summarised for each serum pool and Ig-type (IgG/IgA) in Table 2. Eventually, neighbouring high-fidelity virus-derived peptide pairs were merged into high-fidelity regions (HFR) using the R package IRanges version 2.36.0 [33].

**Table 2.** Number of peptides with high-fidelity antibody binding.

| Organism | Total | Pre-pandemic<br>IgG |  | Pandemic<br>IgG |  | Pre-pandemic<br>IgA | Pandemic<br>IgA |  |
| --- | --- | --- | --- | --- | --- | --- | --- | --- |
| EBOZM | 2417 | 13 (0.54%) |  | 21 (0.87%) |  | 7 (0.29%) | 0 (0.00%) |  |
| HCMV | 31388 | 346 (1.10%) | * | 589 (1.88%) | ** | 38 (0.12%) | 11 (0.04%) |  |
| SARS-CoV-2 | 6330 | 34 (0.54%) |  | 155 (2.45%) | *** | 10 (0.16%) | 41 (0.65%) | ** |
| SARS-CoV | 6353 | 20 (0.31%) |  | 117 (1.84%) | ** | 11 (0.17%) | 27 (0.42%) | ** |
| MERS-CoV | 4933 | 11 (0.22%) |  | 48 (0.97%) |  | 7 (0.14%) | 7 (0.14%) |  |
| HCoV-HKU1(N1) | 4998 | 42 (0.84%) |  | 65 (1.30%) |  | 7 (0.14%) | 5 (0.10%) |  |
| HCoV-OC43 | 5164 | 23 (0.45%) |  | 47 (0.91%) |  | 9 (0.17%) | 5 (0.10%) |  |
| HCoV-NL63 | 4689 | 29 (0.62%) |  | 53 (1.13%) |  | 2 (0.04%) | 4 (0.09%) |  |
| HCoV-229E | 4605 | 25 (0.54%) |  | 53 (1.15%) |  | 5 (0.11%) | 9 (0.20%) |  |

Levels of statistical significance when compared to EBOZM by 2-sample test for equality of proportions with multiple test correction by the Benjamini-Hochberg procedure: \* $p < 0.05$ , \*\* $p < 0.01$ , \*\*\* $p < 0.001$ .
HCMV = human cytomegalovirus, EBOZM = Zaire ebolavirus (strain Mayinga-76), HCoV = Human coronavirus.

The number of HCMV, EBOZM, and HCoV HFRs determined for each serum pool and Ig-type (IgG/IgA) are summarised in Table 3.

**Table 3.** High-fidelity peptides and regions detected in SARS-CoV-2.

| Protein | Total | Pre-pandemic IgA |  |  | Pre-pandemic IgG |  |  | Pandemic IgA |  |  | Pandemic IgG |  |  |
| --- | --- | --- | --- | --- | --- | --- | --- | --- | --- | --- | --- | --- | --- |
|  |  | HFPs | *Fraction | HFRs | HFPs | *Fraction | HFRs | HFPs | *Fraction | HFRs | HFPs | *Fraction | HFRs |
| ORF1ab | 4884 | 5 | 0.10% | 4 | 18 | 0.37% | 11 | 11 | 0.23% | 5 | 37 | 0.76% | 18 |
| S | 636 | 1 | 0.16% | 1 | 11 | 1.73% | 4 | 11 | 1.73% | 6 | 36 | 5.66% | 10 |
| ORF3a | 133 | 0 | 0.00% | 0 | 1 | 0.75% | 1 | 0 | 0.00% | 0 | 10 | 7.52% | 4 |
| M | 131 | 0 | 0.00% | 0 | 0 | 0.00% | 0 | 0 | 0.00% | 0 | 13 | 9.92% | 1 |
| N | 269 | 4 | 1.49% | 1 | 1 | 0.37% | 1 | 19 | 7.06% | 3 | 59 | 2.93% | 7 |
| ORF9b | 47 | 0 | 0.00% | 0 | 3 | 6.38% | 1 | 0 | 0.00% | 0 | 0 | 0.00% | 0 |

\* Fraction indicates the fraction of HFPs out of the total number of peptides investigated for each protein.
HFPs = High-fidelity peptides, HFRs = High-fidelity regions

### Alignment of high-fidelity antibody-binding peptide regions

Unique high-fidelity regions (HFRs) derived from the viruses were uploaded in a single batch to the IEDB Epitope Cluster Analysis web tool (http://tools.iedb.org/cluster/) [34] and clustered against each of the seven HCoV proteomes using a clustering threshold of 60% sequence identity and clustering method 1 (“all connected peptides in a cluster”). The 60% level of homology was empirically selected as the lowest level of homology, which avoided clustering of EBOZM (EP) and HCMV (PP) epitopes with HCoV (CP) epitopes and/or HCoV proteomes (in reality, the 60% sequence identity threshold did inadvertently include a single PP epitope; lower thresholds would have increased these false positive clustering results dramatically). Thus, a clustering threshold of 60% sequence identity would maximise epitope clustering to the HCoV proteomes while avoiding the inclusion of unrelated virus-derived epitopes thereby achieving a reasonable balance between sensitivity and specificity.

The clustered high-fidelity regions were loaded into R [32]. Sequences within each cluster and its subclusters were re-aligned with profile-to-profile alignment using the R package DECIPHER version 2.30.0 [35]. Alignment clusters between coronaviridae and HCMV are summarised in Tables 4 & 5.

### Statistics

In Table 2, alongside the counts of high-fidelity peptides (HFPs), the epitope densities for the EBOZM, HCMV, and each of the HCoV-derived peptides are summarised, calculated as the fraction of HFPs determined in the total number of peptides synthesised for each organism. The EBOZM-derived peptides were included to represent a negative control pathogen, where seroreactivity would be unlikely in Danish human serum pools. Thus, the epitope densities observed in EBOZM represented the apparent false discovery rate of the HFPs determined for each serum pool and Ig-type. We applied the 2-sample test for equality of proportions to determine if the epitope densities of the HCMV-derived positive control peptides (PPs) and the HCoV-derived peptides (CPs) were significantly enriched compared to the serum pool and Ig-type FDRs (i.e. EBOZM epitope densities). P-values derived from the two-proportions z-tests were corrected for multiple testing by the Benjamini-Hochberg procedure, and the resulting significance indicators are summarised in Table 2.

To assess whether the pandemic serum pool pathogen-specific IgG epitope densities were significantly enriched compared to the pathogen-specific IgG epitopes observed for the pre-pandemic serum pool, we applied a one-way exact binomial test. In this test, the number of IgG HFPs in the pandemic serum pool was considered the number of successes, the total count of pathogen-specific peptides served as the number of trials, and the IgG epitope density from the pre-pandemic serum pool was used as the hypothetical probability of success. P-values for each organism-wise comparison were corrected for multiple testing by the Benjamini-Hochberg procedure.

## Results

This study aimed to identify human coronavirus-specific linear B cell epitopes using serum pools collected before or during the early stages of the COVID-19 pandemic. Overlapping peptides representing the complete proteomes of the seven human coronaviruses (HCoVs) were generated by high-density, in situ synthesised peptide microarrays (Table 1). To analyse the binding of IgG and IgA antibodies from the serum pools to the peptides, we first subtracted background signals to account for non-specific binding. For each serum pool and Ig isotype, the background signal distribution was established using randomly generated peptides as negative control peptides (NP). To identify peptides that qualify as high-fidelity linear B cell epitopes, we exploited that our data came from the analysis of overlapping 13-mer peptide libraries with an overlap of 11-amino acids. Since linear B cell epitopes are often shorter than 10 amino acids long, they would tend to appear in at least two consecutive overlapping peptides, appearing as if “stepping” in and out of an immunogenic region with elevated antibody binding. In the Methods section, we described how we harnessed this information about overlapping peptides by integrating the background signal probabilities of a virus-derived peptide with its immediate neighbors, sharing an 11 amino acid overlap in the parent protein, to calculate a combined background probability within the framework of the parent protein sequence. We applied a statistical threshold of a combined p-value ≤ 0.001 to determine peptides likely harbouring linear B cell epitopes. Additionally, to ensure that only peptides eliciting stronger signals were included, we also applied a threshold for antibody binding signal of ≥ 20% of the maximum signal. Peptides satisfying both criteria were categorised as being high-fidelity binding peptides (HFPs). This approach enabled the quantitative assessment of the numbers of pathogen-specific HFPs found before and early into COVID-19 pandemic (Table 2).

The results of this analysis could be visualised by scatter plots (Figure 1 (IgG) and 2 (IgA). To allow for direct comparison of signal values, signals were scaled to the serum pool and Ig-type maximum values and are plotted as the scaled signals from the pandemic serum pools on the Y-axis against the normalised signals from the pre-pandemic serum pools on the X-axis. Whether a given peptide could be categorised as HFP by both the pre-pandemic and pandemic sera, by the pre-pandemic serum only, or by the pandemic serum only, is indicated by the colouring of the figure symbols (Figure 1 (IgG) and 2 (IgA). To facilitate comparisons of immunogenicity across proteomes of different lengths, we have adjusted for protein length by dividing the number of HFP found in each protein with the total number of overlapping peptides representing said proteome (Table 2). Henceforth, we will refer to this length-adjusted measure of epitope recognition as “epitope density”, or simply “densities”.

**Figure 1.**
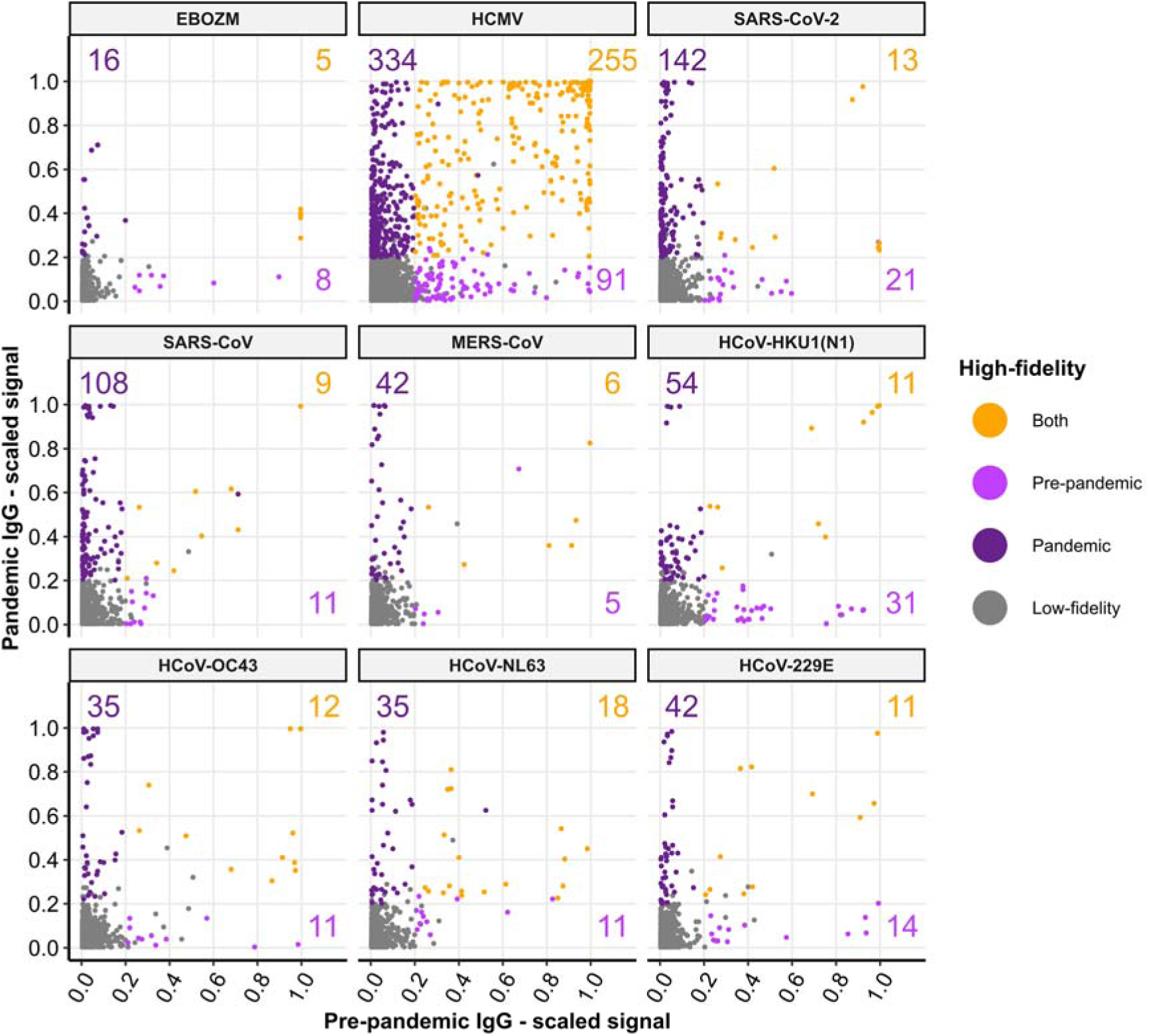
Peptide IgG binding signals on the microarrays. Scatter plot comparing peptide microarray IgG binding signals between the pandemic (y-axis) and pre-pandemic (x-axis) serum pools scaled to the maximum signal measured for each serum pool. Each data point represents one peptide derived from one of the organisms indicated in the plot subtitle. High-fidelity peptides passed our fidelity filters: combined p-value ≤ 0.001 & scaled signal ≥ 20% of the maximum scaled signal measured for the individual serum pool and Ig-type. The legend indicates if a peptide was determined to have high-fidelity IgG binding in the pre-pandemic serum pool (light purple), pandemic serum pool (dark purple), or in both serum pools (orange). Counts indicate the numbers of peptides from each organism determined to have high-fidelity IgG binding distinctly in the pandemic, pre-pandemic, or both serum pools.

As a “non-overlapping” negative control, we used 3900 randomly generated 13-mer peptides (NP). Since these peptides have no overlapping neighbouring peptides, we only applied the “≥ 20% of the maximum signal” criteria. A mere 0.28% of the NP yielded an IgG signal of this magnitude when detected by the pre-pandemic and 0.82% of the NP by the pandemic serum (only 0.10% of the NP were shared). As a more natural “overlapping” negative control, we used 2417 overlapping peptides representing the complete proteome of Zaire ebolavirus (strain Mayinga-76) (EBOZM). Here, 0.54% and 0.87% of the EBOZM-derived peptides (EPs) qualified as HFP when detected by the pre-pandemic and pandemic sera, respectively (only 0.21% of the EP were shared). Using these densities as an approximation, we assume that the IgG HFP false discovery rate (FDR) is 0.54% and 0.87% for the pre-pandemic and pandemic serum pools, respectively (Table 2). By comparison, we found the IgA HFP FDRs to be 0.29% and 0% for the pre-pandemic and pandemic serum pools, respectively (Table 2).

As a positive control (PP), we used 31688 overlapping peptides representing the complete proteome of the HCMV. Given HCMV’s latent nature and its tendency to reactivate periodically, it has typically triggered a robust immune response. Due to its widespread prevalence in the population, we expected that peptides from HCMV would be recognised by serum pools from any randomly selected group of eight individuals. Our results confirmed this expectation: 1.10% and 1.88% of the peptides in the positive control qualified as HFPs when analysed using pre-pandemic and pandemic sera, respectively, with 0.81% being recognised by both. Comparatively, these densities were significantly higher than the corresponding EP densities (P < 0.05 and P < 0.01 for the pre-pandemic and pandemic serum pools, respectively (2-sample test for equality of proportions with multiple test correction by the Benjamini-Hochberg procedure. Hence forward: 2StepBH). As expected, the density of shared pre-pandemic/pandemic HCMV epitopes was the highest of any virus tested here.

### Early COVID-19 pandemic sera exhibited robust IgG and IgA responses against SARS-CoV and SARS-CoV-2

The strategy described above was used to identify high-fidelity linear B cell epitopes from HCoVs. Compared to the IgG and IgA epitope density of EBOZM, the pre-pandemic serum pool HCoV epitope densities were not significantly enriched (Table 2). Reassuringly, this indicated no prior exposure to any of the epi- or pandemic HCoVs in the pre-pandemic serum pool. In contrast, the pandemic serum exhibited significantly higher densities of IgG and IgA epitopes specific to SARS-CoV-2 and SARS-CoV, at 2.45% and 1.84% for IgG, compared to 0.87% for EBOZM IgG, and at 0.65% and 0.42% for IgA, compared to 0% for EBOZM IgA, (P < 0.01, 2StepBH), reflecting the ongoing COVID-19 pandemic. As expected, the SARS-CoV-2 epitope density was the highest of any virus tested in the pandemic serum pool. Finally, neither the pre-pandemic nor the pandemic serum pools had significantly enriched linear B cell epitope densities from any of the common cold coronaviruses (CCCs).

By visual comparison of the response strengths measured as the scaled signals between IgG (Figure 1) and IgA (Figure 2), IgG elicited higher antibody binding signals overall, likely reflecting the serum concentration differences between the two Ig-types. This trend of higher IgG binding translated into the response breadths measured as the numbers of IgG vs IgA HFPs, which were 6- and 10-fold higher in the pre-pandemic and pandemic serum pools, respectively (Table 2). Interestingly, comparing virus-specific IgG epitope densities between the pandemic and pre-pandemic serum pools (Table 2), we found that the pandemic serum pool identified significantly more IgG HFPs across all the investigated viruses (Exact binomial test with multiple test correction by the Benjamini-Hochberg procedure, p-values < 0.01). The underlying mechanisms contributing to this overall enrichment in pandemic IgG responses are not immediately clear.

**Figure 2.**
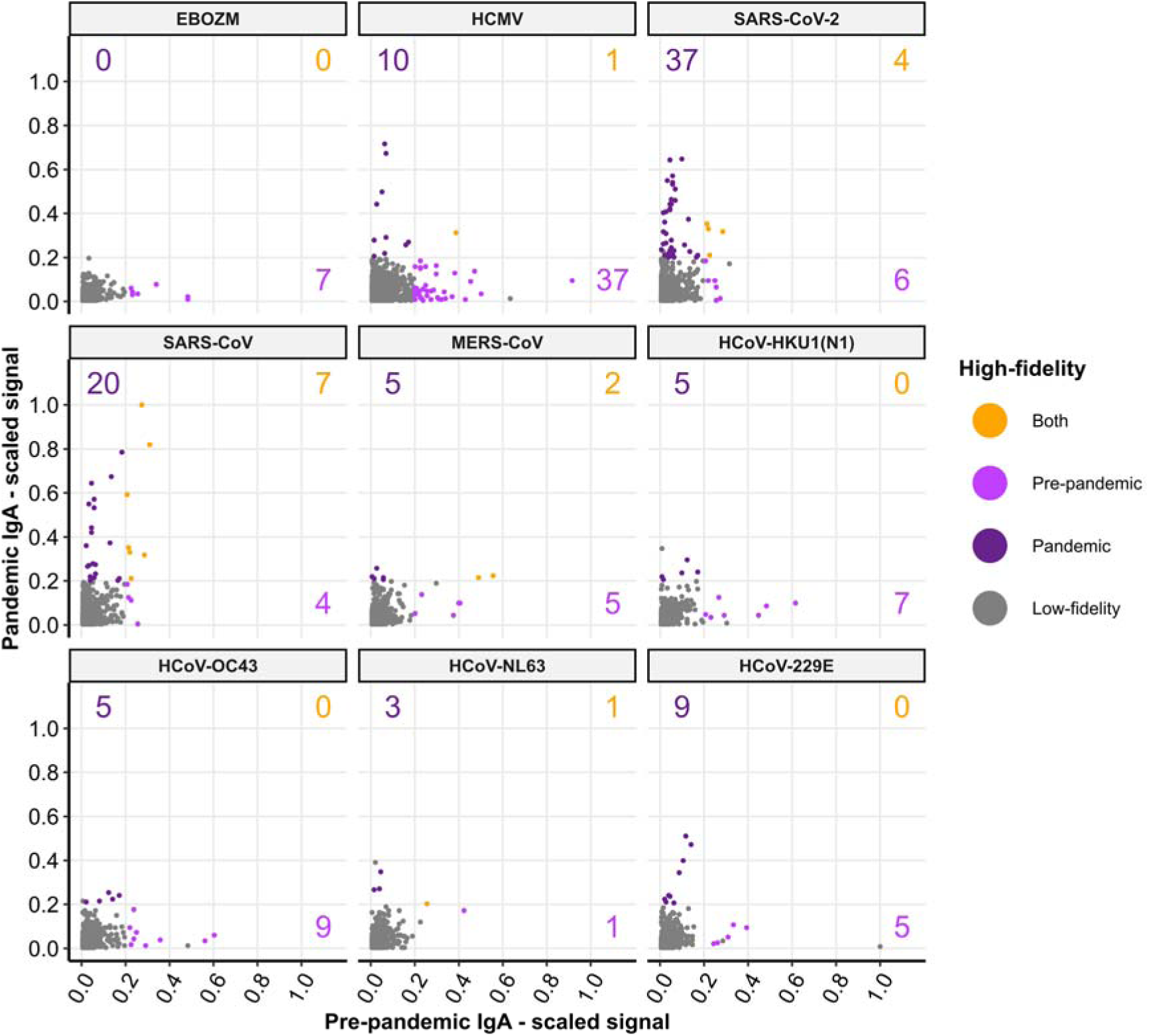
Peptide IgA binding signals on the microarrays. Scatter plot comparing peptide microarray IgA binding signals between the pandemic (y-axis) and pre-pandemic (x-axis) serum pools scaled to the maximum signal measured for each serum pool. Each data point represents one peptide derived from one of the organisms indicated in the plot subtitle. High-fidelity peptides passed our fidelity filters: combined p-value ≤ 0.001 & scaled signal ≥ 20% of the maximum scaled signal measured for the individual serum pool and Ig-type. The legend indicates if a peptide was determined to have high-fidelity IgG binding in the pre-pandemic serum pool (light purple), pandemic serum pool (dark purple), or in both serum pools (orange). Counts indicate the numbers of peptides from each organism determined to have high-fidelity IgA binding distinctly in the pandemic, pre-pandemic, or both serum pools.

### Coronavirus high-fidelity peptides primarily map to the nucleoprotein, the spike protein and the ORF1ab

Focusing on the SARS-2-CoV proteome, we mapped the scaled IgG and IgA binding signals of all the SARS-CoV-2 HFPs to their positions in the proteome (Figure 3). To illustrate this comprehensively, the SARS-CoV-2 proteome is depicted as a central line indicating the size and order of the SARS-CoV-2 proteins and the open reading frames (ORFs) in the proteome. The scaled IgG (blue) and IgA (red) signals of each SARS-CoV-2 HFP are shown as bars with a width corresponding to their length, placed corresponding to their position in the proteome and extending from the centre upwards for the pandemic responses or the centre downwards for the pre-pandemic responses. Non-binding peptides are indicated by grey bars. The highest number of SARS-CoV-2-specific linear B cell epitopes were recognised by IgG in the pandemic serum pool (155 HFPs), followed by IgA in the pandemic serum pool (41 HFPs), IgG in the pre-pandemic pool (34 HFPs) and IgA in the pre-pandemic pool (10 HFPs) (Table 2). When stratifying these HFPs across their SARS-CoV-2 parent proteins, the most immunogenic SARS-CoV-2-derived proteins appeared to be the nucleoprotein (59 IG HFPs and 19 IgA HFPs), followed by the ORF1ab (37 IG HFPs and 11 IgA HFPs) and/or the spike protein (36 IG HFPs and 11 IgA HFPs) recognised by the pandemic serum pool (Table 3). In contrast, the ORF1ab appeared to be the most frequently recognised SARS-CoV-2 protein by the pre-pandemic serum pool (18 IG HFPs and 5 IgA HFPs), probably due to cross-reactions with CCCs (see below).

**Figure 3.**
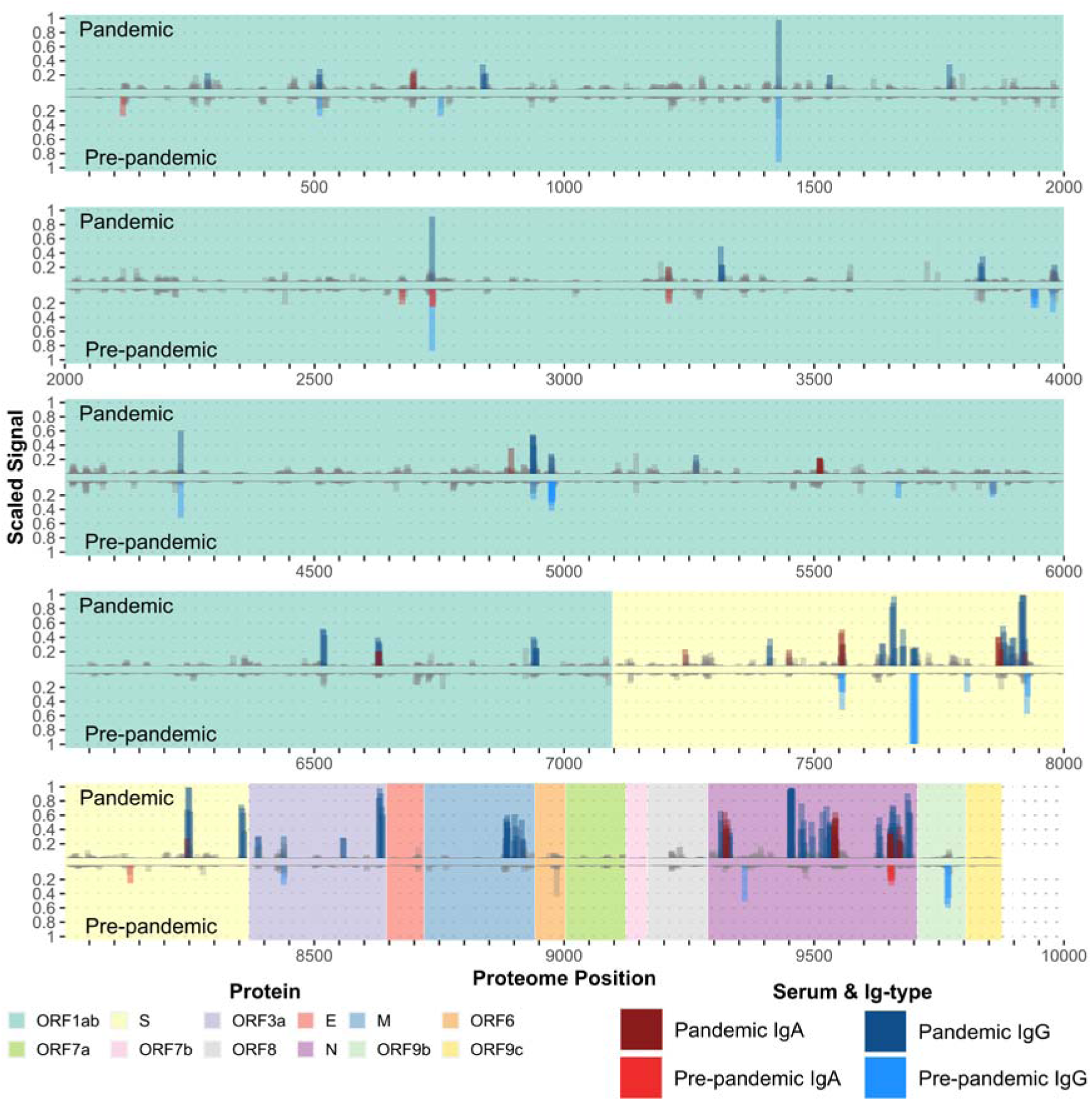
SARS-CoV-2 peptide antibody binding signals mapped to proteome. The y-axis displays standardised IgG and IgA signals on individual microarrays, scaled to their maximum values. Grey bars denote non-binder peptides for both IgG and IgA, while blue/dark blue bars represent high-fidelity IgG-binding peptides (HFPs), and red/dark red bars represent IgA-binding HFPs. Each bar corresponds to the width of a peptide (13 amino acids) and has reduced opacity for overlapping bars. Pandemic serum pool peptide signals are oriented upwards, and pre-pandemic control serum pool peptide signals are oriented downwards. The x-axis shows the SARS-CoV-2 proteome concatenated N- to C-terminal with tick marks at every 50th amino acid position. Each SARS-CoV-2 protein is indicated by the coloured rectangles in the background and is ordered by their start codon position in the reference genome Wuhan-1 (NC_045512.2). HFP = high-fidelity peptide, ORF = Open Reading Frame, S = Spike protein, E = Envelope protein, M = Membrane protein, N = Nucleocapsid protein.

Instead of using the absolute numbers of HFP, one could also use the protein-wise linear B cell epitope densities, thereby adjusting for the differences in lengths and numbers of peptides derived from these proteins. The shortest and longest immunogenic SARS-CoV-2 proteins determined here range from ORF9b at 97 amino acids in length, to ORF1ab, which is 7096 amino acids in length, resulting in the synthesis of 47 and 4884 peptide sequences, respectively (Table 3). By this token, the most densely pandemic IgG HFP populated of the five immunogenic proteins remained the nucleoprotein, and the least is the ORF1ab (out of the proteins with responses, Table 3). These findings concerning which coronavirus proteins were the most immunogenic in terms of absolute numbers and densities of epitopes could be extended to the entire set of HCoV (data not shown).

The HFPs recognised by the pre-pandemic serum pool tended to overlap with those recognised by the pandemic serum pool. This was particularly noteworthy for the ORF1ab. One could speculate whether these are caused by pre-existing antibody responses against endemic CCC and/or cross-reactions between SARS-CoV-2 and CCC responses. In contrast, many of the HFPs in the structural proteins of the SARS-CoV-2 virion particle (Spike (S), Membrane (M), and Nucleocapsid (N) proteins) and ORF3a were exclusive to the pandemic serum pool (Figure 3). Several of the HFPs recognised by the pandemic IgA were found in the S and N proteins overlapping the IgG HFPs, suggesting that these proteins are highly immunogenic antigens capable of eliciting diverse antibody responses across multiple Ig-isotypes.

### Merging high-fidelity peptides into high-fidelity regions (HFR)

Many of the HFPs from the same serum pools and Ig-types overlapped (Figure 3). The exact footprint(s) of the linear B cell epitopes in the overlapping HFPs cannot be determined from these data; however, we reasoned that the footprints could occur within several overlapping HFPs and that neighbouring and overlapping footprints could merge into high-fidelity regions (HFRs). Such an approach could encompass larger immunogenic structures. Overall, the average length of these HFRs was 18 amino acids, representing 3-4 merged HFPs (range 13 to 157 amino acids, representing 1 to 19 HFPs). The number of unique HFRs detected in each organism, serum pool, and Ig-type grouping are summarised in Table 4. In total, we identified 280 unique HCoV HFRs, 201 HFRs by the pandemic serum pool and 102 by the pre-pandemic pool (53 HFRs were identical between the two serum pools). As expected, and echoing the HFP analysis of HCoV above, the largest contribution, with 72 HFRs, came from SARS-CoV-2, the virus that caused the ongoing pandemic. The second largest contribution, with 53 HFRs, came from the closely related SARS-CoV, likely reflecting cross-reactions from SARS-CoV-2.

**Table 4.** High-fidelity regions detected in human coronaviruses, human cytomegalovirus, and ebolavirus.

| Organism | Pre-pandemic IgA | Pre-pandemic IgG | Pandemic IgG | Pandemic IgA | Organism union* |
| --- | --- | --- | --- | --- | --- |
| HCMV | 21 | 96 | 165 | 7 | 271 |
| EBOZM | 5 | 6 | 11 | 0 | 21 |
| SARS-CoV-2 | 6 | 18 | 40 | 14 | 72 |
| SARS-CoV | 6 | 10 | 32 | 9 | 53 |
| HCoV-HKU1(N1) | 4 | 17 | 25 | 3 | 47 |
| HCoV-NL63 | 1 | 15 | 24 | 2 | 37 |
| HCoV-229E | 3 | 9 | 21 | 4 | 34 |
| HCoV-OC43 | 5 | 11 | 16 | 4 | 34 |
| MERS-CoV | 4 | 7 | 19 | 4 | 32 |
| HCoVs union** | 26 | 76 | 167 | 34 | ***280 |

\* Organism total shows the unique high-fidelity region (HFR) sequences detected across all serum pools and Ig-types for the indicated organisms.
\*\* HCoVs total shows the unique HFR sequences in each serum pool and Ig-type across all HCoVs.
\*\*\* The number of unique HFR sequences found across all HCoVs, serum pools, and Ig-types.
HCMV = human cytomegalovirus, EBOZM = Zaire ebolavirus (strain Mayinga-76), HCoV = Human coronavirus.

### Homology clustering of high-fidelity regions suggests antibody cross-reactivity between human coronaviruses

To assess the potential of antibody cross-reactivity against the HCoV-derived HFRs, all the unique HFRs identified from the seven HCoVs, HCMV and EBOZM, were submitted to the Immune Epitope Database (IEDB) Epitope Cluster Analysis web tool (http://tools.iedb.org/cluster/). To avoid false positive clustering, we empirically selected a sequence identity threshold of 60% where only 1 of the 271 HCMV HFRs and none of the 21 Ebola HFR clustered with the HCoV HFRs. The alignments and clustering of all the HCoV HFRs are summarised in Supplementary Table 1. For completeness, the alignments and clustering of all HCMV and EBOZM HFRs are summarised in Supplementary Table 2.

Of the 333 (280 unique) HCoV-derived HFRs, 225 HFRs could be grouped into 78 unique clusters of 2-15 HFR members (Table 3). The remaining 108 HFRs were singletons, which did not cluster with any other HCoV HFRs. Each of the 186 clusters and singletons was given a unique Alignment ID number from 1 to 186. Figure 4A features a 4-circle Venn diagram that categorises alignment clusters as either shared or uniquely recognised, contingent upon the 4 combinations of serum pools: pre-pandemic or pandemic and IgG or IgA (detailed in Figure 4B). Notably, 138 out of 186 (74%) of these clusters were exclusive to specific serum and isotype pairings (Figure 4A). Conversely, shared clusters among two, three, and all four combinations were less common, with counts of 41 (22%), 7 (4%), and 0 (0%), respectively (Figure 4A). This relative lack of shared recognition underscores the previously proposed individuality of linear B cell recognition. From a more practical point of view, searching for possible cross-reactions against different HCoV strains becomes more challenging simply because there are fewer examples of shared recognition to investigate.

**Figure 4.**
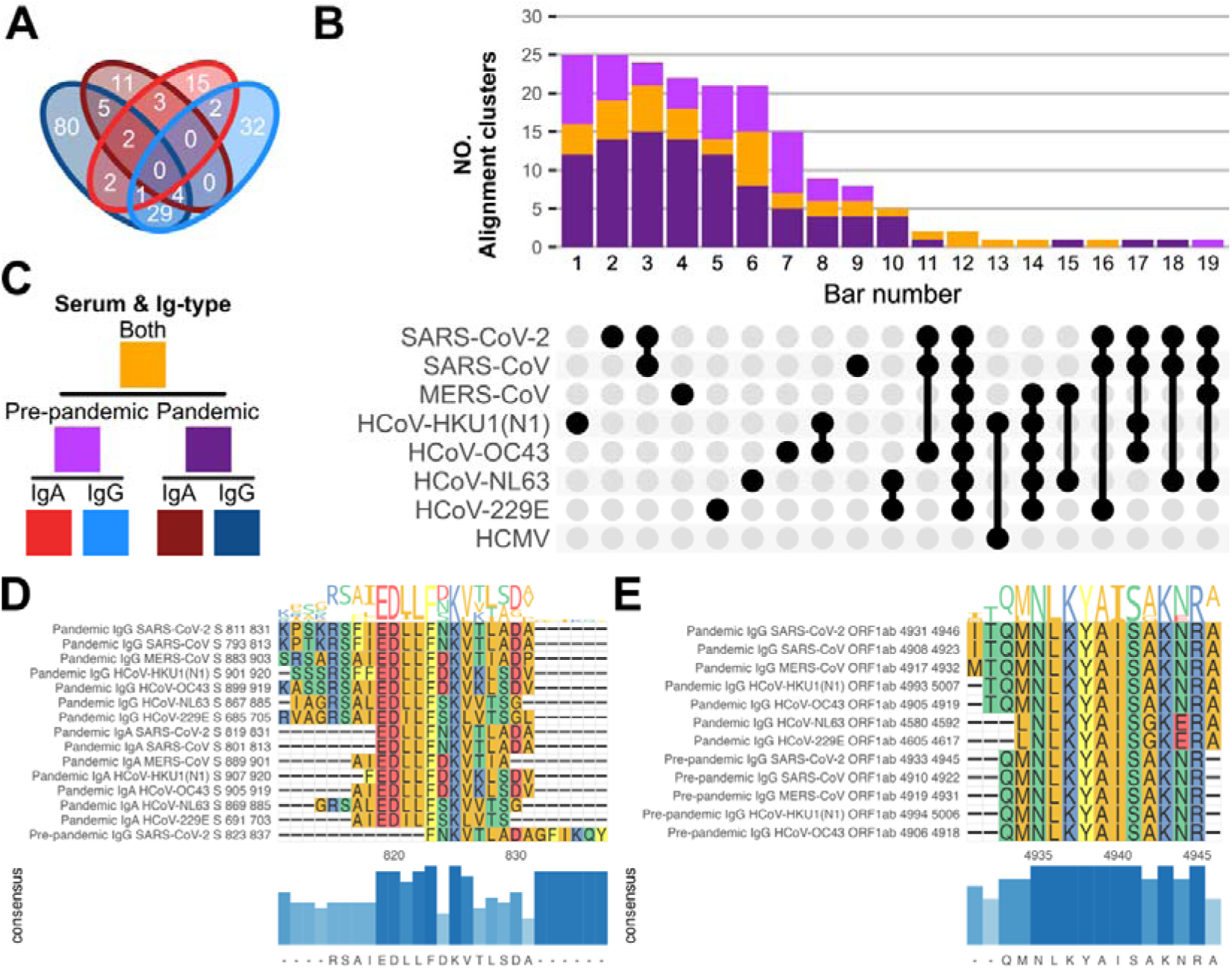
Summary of high-fidelity region alignment clusters. The 333 (280 unique) high-fidelity regions (HFRs) from both serum pools and Ig-types derived from the human coronaviruses (HCoVs) (Table 2) were clustered into 186 alignment clusters using single-linkage clustering with a sequence similarity cut-off at 60% (Supplementary Table 1). **A**: Venn diagram showing how the 186 alignment clusters are distributed or shared between the serum pool and Ig-types. **B**: Stacked bar plot with counts of how many alignment clusters are specific to the pandemic serum, the pre-pandemic control serum, or if the alignment clusters have HFRs from both sera. The combination matrix below indicates which organisms were aligned. **C**: Is a pedigree chart legend indicating how colours relate to each other in **A** and **B**. **D**: Shows the aligned HFRs with alignment ID 135 (Supplementary Table 1). The alignment covers positions 811 to 837 in the SARS-CoV-2 Spike (S) protein. **E**: Shows the aligned HFRs with alignment ID 87 (Supplementary Table 1). The alignment covers positions 4931 to 4947 in the SARS-CoV-2 Open Reading Frame (ORF) 1ab protein. To the left of the aligned HFRs (**DE**) are their serum pools, Ig-types, organisms, parent proteins, and start/end positions. Above, sequence logos represent amino acid frequencies per position. Amino acids are color-coded by sidechain chemistry. Below, a bar plot shows each position’s most frequent amino acid (or gap). The inferred consensus sequence is at the bottom. HFR = high-fidelity region, HCoV = human coronavirus, HCMV = human cytomegalovirus, NO. = number of, S = Spike protein, ORF = Open Reading Frame.

SARS-CoV, MERS-CoV, and SARS-CoV-2 (before the COVID-19 pandemic) have not circulated in Denmark and should only be recognised by the pre-pandemic serum pool through cross-reactions with common cold coronaviruses (CCCs). Likewise, for the pandemic serum pool, antibody responses targeting SARS-CoV or MERS-CoV should be the result(s) of cross-reactivities with either the CCCs and/or SARS-CoV-2. Using the homology clustering approach (Supplementary Table 1), we found that 100% of the IgA HFRs and 81% of the IgG HFRs found in the pandemic serum pool could be explained by cross-reactivity to either SARS-CoV-2 and/or one or more of CCCs, consistent with the sequence similarity between SARS-CoV and SARS-CoV-2. However, for MERS-CoV, only 25% and 21% of the IgA and IgG HFRs could be explained by alignment to SARS-CoV-2 and/or the CCCs. This lower degree of alignment is consistent with MERS-CoV sharing the least sequence similarity with the other HCoVs but begs the question of how these antibody responses arose. For the pre-pandemic serum pool, none of the SARS-CoV, MERS-CoV, or SARS-CoV-2 HFRs aligned with any of the CCCs. For the pre-pandemic IgG HFRs, 30%, 29%, and 17% of the responses against SARS-CoV, MERS-CoV, or SARS-CoV-2 could be explained by alignment with one or more of the CCC-derived HFRs, respectively. The presence of unexplained pre-pandemic antibody recognition of SARS-CoV, MERS-CoV, and SARS-CoV-2 derived epitopes suggests that further improvements of the present approach is warranted (see the discussion).

A more detailed analysis of HCoV cross-reactions is summarised in Figure 4C. Alignment clusters were simplified by merging IgG or IgA HFRs, as illustrated in the pedigree tree in Figure 4B. The 19 numbered bars represent the counts of alignment clusters with HFRs from different combinations of HCoVs (Figure 4B). Whether the members of these alignment clusters were recognised by the pre-pandemic or pandemic serum pools or by both is indicated by the segmental colouring of the bars. Alignment cluster bar numbers are cross-referenced in Supplementary Table 1. By qualitative assessment of the bars (Figure 4B), four main groups were identified: (I) the epi- and pandemic HCoVs of the betacorona genus, SARS-CoV, MERS-CoV, and SARS-CoV-2, represented by bars 2, 3, 4, and 9, which cover the majority of alignment clusters and were predominantly recognised by pandemic serum pool; (II) the CCCs of the alphacorona genus (HCoV-NL63 and HCoV-229E), represented by bars 5, 6, and 10, which were more evenly distributed between the serum pools; (III) the CCCs of the betacorona genus, HCoV-HKU1(N1) and HCoV-OC43, represented by bars 1, 7, and 8, which were also more evenly recognised by the pre-pandemic and pandemic serum; and (IV) a pan-HCoV specific group represented by bar 12, which happens to consist of the two single largest alignment clusters (Supplementary Table 1, Align ID 87 & 135), one of 15 members from the S protein (Figure 4D) and another of 12 members from the ORF1ab polyprotein (Figure 4E), which were shared between the pre-pandemic and pandemic serum pools. Finally, eight smaller bars (Figure 4B, bar numbers 11, 13-19) represented alignment clusters with HFRs from various HCoVs across genera.

Figure 4D displays the largest alignment cluster (Align ID 135, Supplementary Table 1). 14 of 15 HFRs were recognised by the pandemic serum pool (half by IgG, half by IgA); only one HFR was recognised by the pre-pandemic serum pool IgG, and since it was shifted C-terminally compared to all the other members of this alignment group, one could argue that it might not belong to the alignment. Thus, a consensus sequence consisting of 815-RSAIEDLL-822 is likely to be a dominant, pan-HCoV-specific linear B cell epitope footprint elicited by the SARS-CoV-2 infection.

Figure 4E, displays the second largest alignment cluster (Align ID 87, Table 3) located between positions 4931 and 4947 when mapped onto the SARS-CoV-2 ORF1ab protein. Five of the 12 HFRs were recognised by pre-pandemic serum pool IgG and the remaining seven were recognised by pandemic serum pool IgG. This linear B cell epitope cluster exemplifies a pan-HCoV specific, encompassing all seven HCoV strains.

## Discussion

We analysed linear B cell epitopes from SARS-CoV-2 and 6 other pandemic or epidemic human coronavirus strains. We obtained pooled sera from COVID-19 convalescents (i.e. SARS-CoV-2 exposed) sampled early in the pandemic (around March 2020) and from pre-pandemic (not SARS-CoV-2 exposed) individuals sampled before October 2019 and tested them for relevant IgG and IgA responses. To detect linear B cell epitopes, high-density peptide microarrays were used to generate overlapping 13 amino acid peptides tiling by 2 amino acids to represent the entire proteomes of all seven HCoVs. These microarrays were incubated with pre-pandemic or pandemic serum pools, and IgG and IgA reactivity against specific peptides were detected. We identified 280 regions of linear B cell epitopes recognised by the pre-pandemic and/or pandemic serum pools. These regions could be homology clustered and aligned to the HCoV proteomes, providing evidence of cross-reactions between the pandemic and epidemic HCoVs. These clusters might be of therapeutic and diagnostic value: the largest cluster was pan-HCoV specific and has, by others, been implicated in the neutralising activity of sera from COVID-19 convalescents [36–40], the second largest cluster appeared to be pan-HCoV-specific, whereas other smaller clusters appeared to be strain or group specific.

Here, we have identified a large cohort of SARS-CoV-2 and HCoV-derived linear B cell epitopes and generated data on the IgG and IgA-mediated humoral responses towards these epitopes. At the protein level, we show that the humoral immune system primarily targets the ORF1ab protein, the Spike protein, and the nucleoprotein, with minor targets including ORF3a and ORF9b. Others have obtained similar results studying antibody responses against SARS-CoV-2 proteins after vaccination or natural infection. Currently (November 2023), 8,528 linear B-cell epitopes have been submitted to the Immune Epitope Database (IEDB) [44]. The majority of the methods used for linear antibody epitope discovery in SARS-CoV-2 include enzyme-linked immunosorbent assay (ELISA) [36–38,45–67], peptide arrays [68,69], peptide microarrays [70–84], phage display [43,85–93], or bacterial display [94]. The top three SARS-CoV-2 source proteins of linear B cell epitopes in the IEDB data are the ORF1ab (54%), S (30%), and N (9%) proteins [44].

When trying to understand the interactions between a virus and its host cells that lead to infection, the value of linear B cell epitope information may be questioned. In the case of the SARS-CoV-2 infection, the infective event involves structural rearrangements of the spike protein of the virus (reviewed in [41]), and short synthetic peptides are not likely to mimic faithfully structural events involving larger, often discontinuous, viral structures.

Nonetheless, linear information might be quite valuable. Poh et al. 2020 identified two peptides that successfully adsorbed neutralising activity from the convalescent sera [36]. One of them, S21P2, was contained in the largest cluster found in our study, Alignment ID 135 (SARS-CoV-2 spike protein 811-831, K<u>PSKRSFIEDLLFNKV</u>TLADA, underscoring indicating the rendition of this epitope reported by Poh et al. 2020) and the other, S14P5, was identified as a singleton in our study, Alignment ID 131 (SARS-CoV-2 spike protein 553-571, <u>TESNKKFLPFQQFGRDIA</u>D, underscoring indicating the rendition of this epitope reported by Poh et al. 2020). Low et al. 2022 isolated monoclonal antibodies specific for SARS-CoV-2 spike protein from convalescent donors [40]. Seven of 16 of these recognised a peptide that was also contained in our Alignment ID 135 (SARS-CoV-2 spike protein 811-831, K<u>PSKRSFIEDLLFNK</u>VTLADA; underscoring indicating the rendition of this epitope reported by Low et al. 2022). Two of these seven monoclonal antibodies showed pan-HCoV-specific neutralisation in vitro and protection in vivo using animal challenge studies. Linear B cell epitopes might also be of diagnostic value. Shrock et al. 2020 used a bacteriophage library approach (“VirScan” [42]) expressing 56-mer peptides tiling by 28 amino acids representing proteomes of all known pathogenic human viruses to examine the serological response against HCoVs and identify linear B cell epitopes [43]. Using this data, they developed a machine learning model that could accurately distinguish between COVID-19 positive and negative individuals.

Apart from the therapeutic and diagnostic implications described above, linear B cell epitope information might also contribute to explaining the natural course of COVID-19. Specifically, the potential cross-reactivity between the pandemic SARS-CoV-2 and endemic CCCs may indicate the presence of pre-existing immunity, offering a possible explanation for the highly variable outcome of the infection. To investigate the possibility of cross-reactions among linear B cell epitopes from different HCoV origins, we conducted a systematic comparison of all identified linear B cell epitopes and asked specifically whether epitope similarities could account for the observed humoral immune responses against SARS-CoV-1 and MERS, despite these viruses not having circulated in Denmark. Since many of the SARS-CoV-2 HFPs in Figure 3 overlapped, we initially merged the overlapping HFPs to reduce the HFPs into high-fidelity regions (HFRs) covering the linear B cell epitope footprints. While the average length of these HFRs was 18 amino acids, representing 3-4 merged HFPs, certain HFRs extended up to 157 amino acids, potentially encompassing multiple linear B cell epitopes simultaneously. We then used the Immune Epitope Database (IEDB) Epitope Cluster Analysis web tool (http://tools.iedb.org/cluster/) [34] to homology cluster HFRs based on a 60% sequence identity threshold. This threshold was selected to avoid false positive clustering of linear B cell epitopes from unrelated organisms; *in casu*, from HCMV. Using this threshold, we identified 78 homology clustered alignments between the HCoVs. Some of the HFRs from SARS-CoV and MERS-CoV occurred in clusters with CCCs and/or SARS-CoV-2, which could explain why they were recognised by the pre-pandemic and/or pandemic sera, respectively. Our data adds to the notion that pre-existing immunity to the endemic CCCs could cross-react with the epi- and pandemic HCoVs. This could lead to partial or complete protection depending on the unknown individual history of CCC exposure; something that for practical purposes is an unpredictable and confounding prognostic factor. Additionally, we observed many homology clusters between SARS-CoV and SARS-CoV-2 (Figure 4B, Bar number 3), which could explain SARS-CoV-1 responses in the pandemic serum pool. The latter potential for antibody cross-reactivity between SARS-CoV and SARS-CoV-2 is well described [43,61,69,72,73,77,85,94–99], and some of these antibodies have been shown to lead to cross-neutralization between these two closely related HCoVs [37,39,40,52,57,59,64,67,70,79,81,100–102]. When this is said, homology clustering did not account for all unexpected seroreactivities, as both serum pools retained SARS-CoV- and MERS-CoV-specific antibody responses. Additionally, the negative serum exhibited SARS-CoV-2-specific reactivities that did not align with any of the CCCs. The unexplained exclusive HFRs may result from cross-reactive linear B cell epitope footprints below the homology threshold. The alphacorona HCoVs and MERS-CoV overall share less than 30% protein sequence identity in the S protein with the betacorona HCoVs [6]. Alternatively, cross-reactivity to past microorganism exposures not addressed in this study could be at play. For instance, the SARS-CoV-2 S protein S2 subunit (S686-1273) has been reported to harbour cross-reactive epitopes elicited by commensal microorganisms [103]. Finally, homology clustering can only suggest cross-reactivity between the different HCoV HFRs. Alanine scans or complete substitution analyses must confirm cross-reactivity to elucidate the linear B cell epitope footprint positions and amino acid specificities.

An interesting observation was a significant increase in the number of HCMV and Zaire Ebola virus responses in the pandemic vs pre-pandemic serum pools. A priori, one would not expect that an infection with SARS-CoV-2 would increase humoral responses to the unrelated HCMV and Ebola virus. Shrock et al. 2020 [43] also reported increased seroreactivity against HCMV and herpes simplex virus 1 in non-hospitalized COVID-19 patients compared to hospitalised. We do not know the reason for this increased humoral reactivity in COVID-19 convalescents. Others have suggested that SARS-CoV-2 relaxes peripheral B cell tolerance [104] and/or causes polyclonal “by-stander” B cell activation [105]. Notably, Jiang et al. 2023 also reported significantly increased anti-HCMV IgG titers in their COVID-19 patient sera [105].

Finally, we would like to discuss the technological implementation of this technology and possible improvements. A major advantage of addressing linear B cell epitopes as opposed to conformational B cell epitopes lies in the exhaustive nature and relative ease of a linear peptide-driven analysis. Peptides, the embodiment of linear B cell epitopes, can be procured as synthetic high-density peptide microarray libraries featuring hundreds of thousands of addressable peptides, enough to encompass multiple virus proteomes. Which sequences should be used for effort can be determined from DNA sequencing the genomes of interest; for COVID-19, the first complete genome and its proteome derivative became available a few days into the pandemic. Once a peptide has been identified as a linear B cell epitope, the detailed specificity of the (even polyclonal) antibodies recognising this epitope can be virtually atomised using a complete single substitution analysis and, if need be, a multi-substitution analysis. We have demonstrated the power of a complete single substitution analysis (as opposed to a simpler alanine scanning analysis) [27–29] and proposed that an ANOVA analysis can be used to detail this specific in objective, statistical terms (location and spacing of important residues that positively or negatively affects recognition, and identification of such residues). However, for a 15 (or 13-mer as in this analysis) peptide, this would require that 15 x 20 = 300 peptides (or 260 peptides for a 13-mer) be synthesised for each tiling peptide in an overlapping peptide library. We conducted a complete analysis of linear B cell epitopes in human serum albumin along these lines and used about 180,000 peptides to do this. It would be untenable to do this as a first choice for any proteome – even using high-density peptide microarrays as we did here. To solve this conundrum, we propose using neighbouring peptide signals to down-size this problem. In our past large-scale linear B cell epitope screens, where we characterised linear B cell epitopes using a complete single substitution analysis, we observed that most epitopes were sufficiently short to appear in at least two neighbouring, overlapping peptides [27,28]. Thus, we here introduced a criterium requiring peptides for further analysis to be found in at least one neighbouring peptide. We implemented this criterion by calculating the combined probability of both members of any pair belonging to the background population and required this to be ≤ 0.001. To prepare our data for substitution analysis, we further introduced another criterion that also had to be met, having an antibody binding signal of ≥ 20% of the highest antibody binding signal recorded on the microarray. This ensures that any peptide selected for further study can be examined by substitution analysis, which requires that changes in the signals obtained with analogue peptides could clearly be distinguished from the proband peptide and, ideally, also from the background. With these combined requirements, we aim to include most actual linear B cell epitopes in our analysis while avoiding those that a priori are unlikely to be such epitopes and/or be subject to substitution analysis. Here, we use this double requirement to condense individual peptide signals into a binary distinction: peptides with high-fidelity antibody binding (HFPs) meeting both requirements or non-binders. We want to add that these simple requirements would support a two-tiered linear B cell epitope discovery process where an initial scan using overlapping 13-mer peptides tiling by two amino acids could address entire virus proteomes and identify putative linear B cell epitopes for subsequent detailed substitution analysis (unfortunately, we did not have the resources to do the latter in this study). Such substitution analyses would identify peptides motifs consisting of a variable number of preferred (and/or disliked) amino acids in an appropriate arrangement and spacing. This information would provide a detailed and hig-resolution characterisation of each recognised linear B cell epitope and could be used to improve future searches for potentially cross-reaction epitopes.

Another issue for future improvements is the source of the antibodies used in this analysis. COVID-19 spread rapidly in the early days of the pandemic, reaching thousands of cases by January 2020, tens of thousands by early February, and over a hundred thousand cases by early March 2020 [106]. Thus, large numbers of relevant pandemic serums have easily been obtained from convalescent donors. Likewise, relevant pre-pandemic serum could be sourced from suitable existing biobanks. We suggest that the initial scan always be conducted with pools of relevant serum. We have in a previous study, where individual sera were analysed, observed that the linear B cell repertoire is individualised to the extent that very few shared (or public) linear B cell epitopes exists [107]. Others pursuing similar large-scale, linear B cell epitope discovery using brute-force approaches of testing hundreds of individual sera [43] have demonstrated that very few public epitopes, that are recognised in 30-80% of individuals, exist. In contrast, many private epitopes, recognised in less than 5% of the population, exist [43,108]. Here, we found a mere 30% overlap in HCMV reactivity between the pandemic and pre-pandemic serum pools, which suggests some degree of individualisation of linear B cell responses in our study also (Table 4). Such epitope individualisation would seriously impact epitope discovery if identifying public epitopes is important. We interpret this as another indication of the individualised nature of linear B cell epitope recognition. The serum pools used here were only comprised of eight individuals. We suggest that the largest possible pool be used for the initial scan and any follow-up substitution analysis. Everything else being equal, this should focus the analysis on public epitope, which might be of more general utility. Eventually, when linear B cell epitopes of interest have been identified and characterised, then individual sera can be examined using single peptides (rather than entire peptide microarrays) to define the prevalence of linear B cell epitopes in the population.

Overall, the approach described here allows for a fast response to future pandemics and yields valuable data on linear B cell epitopes. Identifying virus-specific B cell epitopes is important for rational approaches to prevent, treat, diagnose, and understand virus infections.

## Conclusion

We provide proof-of-concept for using serum pools on high-density peptide microarrays in linear B-cell epitope discovery from available genetic information. Even with a small pool of relevant pre-pandemic and pandemic sera, our findings correspond with literature regarding antibody reactivity against SARS-CoV-2 and other related human coronaviruses in sera of COVID-19 convalescents and pre-pandemic donors. Homology-clustering of our antibody targets revealed intriguing patterns of potential antibody cross-reactivity between the HCoVs, corroborated in the literature. These findings underscore the complexity of antibody responses and suggested solutions to address cross-reactivity patterns. The risk of future pandemics is substantial given our globalised lifestyle. We propose that the approach taken here, could be a valuable tool in the response to future pandemics allowing the discovery of targets for rational vaccine designs and serological testing.

## Supporting information

Supplementary Table 1. Homology clustering alignments of coronaviridae high-fidelity regions

Supplementary Table 2. Homology clustering alignments of human cytomegalovirus and ebolavirus high-fidelity regions

## Data Availability

The datasets presented in this study can be found in online repositories. The name of the repository/repositories and accession number(s) can be found below: https://doi.org/10.5061/dryad.s1rn8pkg7

## Acknowledgement

This work was supported by the European Commission (Electromed, project id. 862539).

## References

1. Johns Hopkins Center for Systems Science and Engineering (CSSE). COVID-19 Map - Johns Hopkins Coronavirus Resource Center. Published October 31, 2023. Accessed 10, 2023. https://coronavirus.jhu.edu/map.html

2. Chen L, Liu W, Zhang Q, et al. RNA based mNGS approach identifies a novel human coronavirus from two individual pneumonia cases in 2019 Wuhan outbreak. Emerg Microbes Infect. 2020;9(1):313–319. doi:10.1080/22221751.2020.1725399

3. Su S, Wong G, Shi W, et al. Epidemiology, Genetic Recombination, and Pathogenesis of Coronaviruses. Trends Microbiol. 2016;24(6):490-502. doi:10.1016/j.tim.2016.03.003

4. Gussow AB, Auslander N, Faure G, Wolf YI, Zhang F, Koonin EV. Genomic determinants of pathogenicity in SARS-CoV-2 and other human coronaviruses. Proc Natl Acad Sci U S A. 2020;117(26):15193–15199. doi:10.1073/pnas.2008176117

5. Kesheh MM, Hosseini P, Soltani S, Zandi M. An overview on the seven pathogenic human coronaviruses. Rev Med Virol. 2022;32(2):e2282. doi:10.1002/rmv.2282

6. Cicaloni V, Costanti F, Pasqui A, Bianchini M, Niccolai N, Bongini P. A Bioinformatics Approach to Investigate Structural and Non-Structural Proteins in Human Coronaviruses. Front Genet. 2022;13:891418. doi:10.3389/fgene.2022.891418

7. Chen B, Tian EK, He B, et al. Overview of lethal human coronaviruses. Signal Transduct Target Ther. 2020;5(1):89. doi:10.1038/s41392-020-0190-2

8. World Health Organization. Status of COVID-19 Vaccines within WHO EUL/PQ evaluation process. Published August 8, 2023. Accessed February 11, 2023. https://extranet.who.int/pqweb/key-resources/documents/status-covid-19-vaccines-within-who-eulpq-evaluation-process

9. World Health Organization. COVID-19 vaccine tracker and landscape. Published March 30, 2023. Accessed November 2, 2023. https://www.who.int/publications/m/item/draft-landscape-of-covid-19-candidate-vaccines

10. Carabelli AM, Peacock TP, Thorne LG, et al. SARS-CoV-2 variant biology: immune escape, transmission and fitness. Nat Rev Microbiol. 2023;21(3):162–177. doi:10.1038/s41579-022-00841-7

11. Yamasoba D, Kimura I, Nasser H, et al. Virological characteristics of the SARS-CoV-2 Omicron BA.2 spike. Cell. 2022;185(12):2103-2115 e19. doi:10.1016/j.cell.2022.04.035

12. Cao Y, Wang J, Jian F, et al. Omicron escapes the majority of existing SARS-CoV-2 neutralizing antibodies. Nature. 2022;602(7898):657-663. doi:10.1038/s41586-021-04385-3

13. Omotuyi O, Olubiyi O, Nash O, et al. SARS-CoV-2 Omicron spike glycoprotein receptor binding domain exhibits super-binder ability with ACE2 but not convalescent monoclonal antibody. Comput Biol Med. 2022;142:105226. doi:10.1016/j.compbiomed.2022.105226

14. Nextstrain team. Genomic epidemiology of SARS-CoV-2 with subsampling focused globally over the past 6 months. Published November 2, 2023. Accessed November 3, 2023. https://nextstrain.org/ncov/gisaid/global/6m

15. Cao Y, Jian F, Wang J, et al. Imprinted SARS-CoV-2 humoral immunity induces convergent Omicron RBD evolution. Nature. 2023;614(7948):521-529. doi:10.1038/s41586-022-05644-7

16. Faraone JN, Qu P, Goodarzi N, et al. Immune Evasion and Membrane Fusion of SARS-CoV-2 XBB Subvariants EG.5.1 and XBB.2.3. bioRxiv. Published online August 30, 2023:2023.08.30.555188. doi:10.1101/2023.08.30.555188

17. Wang Q, Guo Y, Zhang RM, et al. Antibody neutralisation of emerging SARS-CoV-2 subvariants: EG.5.1 and XBC.1.6. Lancet Infect Dis. 2023;23(10):e397–e398. doi:10.1016/S1473-3099(23)00555-8

18. Lasrado N, Collier ARY, Hachmann NP, et al. Neutralization Escape by SARS-CoV-2 Omicron Subvariant BA.2.86. bioRxiv. Published online September 5, 2023:2023.09.04.556272. doi:10.1101/2023.09.04.556272

19. Hu Y, Zou J, Kurhade C, et al. Less neutralization evasion of SARS-CoV-2 BA.2.86 than XBB sublineages and CH.1.1. bioRxiv. Published online September 11, 2023:2023.09.10.557047. doi:10.1101/2023.09.10.557047

20. Ferdous S, Kelm S, Baker TS, Shi J, Martin ACR. B-cell epitopes: Discontinuity and conformational analysis. Mol Immunol. 2019;114:643–650. doi:10.1016/j.molimm.2019.09.014

21. Sharon J, Rynkiewicz MJ, Lu Z, Yang CY. Discovery of protective B-cell epitopes for development of antimicrobial vaccines and antibody therapeutics. Immunology. 2014;142(1):1–23. doi:10.1111/imm.12213

22. Haste Andersen P, Nielsen M, Lund O. Prediction of residues in discontinuous B-cell epitopes using protein 3D structures. Protein Sci. 2006;15(11):2558–2567. doi:10.1110/ps.062405906

23. Kringelum JV, Lundegaard C, Lund O, Nielsen M. Reliable B cell epitope predictions: impacts of method development and improved benchmarking. PLoS Comput Biol. 2012;8(12):e1002829. doi:10.1371/journal.pcbi.1002829

24. Jespersen MC, Mahajan S, Peters B, Nielsen M, Marcatili P. Antibody specific B-cell Epitope predictions: Leveraging information from antibody-antigen protein complexes. Front Immunol. 2019;10:298. doi:10.3389/fimmu.2019.00298

25. Jespersen MC, Peters B, Nielsen M, Marcatili P. BepiPred-2.0: improving sequence-based B-cell epitope prediction using conformational epitopes. Nucleic Acids Res. 2017;45(W1):W24–W29. doi:10.1093/nar/gkx346

26. Cia G, Pucci F, Rooman M. Critical review of conformational B-cell epitope prediction methods. Brief Bioinform. 2023;24(1). doi:10.1093/bib/bbac567

27. Buus S, Rockberg J, Forsstrom B, Nilsson P, Uhlen M, Schafer-Nielsen C. High-resolution mapping of linear antibody epitopes using ultrahigh-density peptide microarrays. Mol Cell Proteomics. 2012;11(12):1790–1800. doi:10.1074/mcp.M112.020800

28. Hansen LB, Buus S, Schafer-Nielsen C. Identification and mapping of linear antibody epitopes in human serum albumin using high-density Peptide arrays. PLoS One. 2013;8(7):e68902. doi:10.1371/journal.pone.0068902

29. Hansen CS, Østerbye T, Marcatili P, Lund O, Buus S, Nielsen M. ArrayPitope: Automated analysis of amino acid substitutions for peptide microarray-based antibody Epitope mapping. PLoS One. 2017;12(1):e0168453. doi:10.1371/journal.pone.0168453

30. Lachmann R, Loenenbach A, Waterboer T, et al. Cytomegalovirus (CMV) seroprevalence in the adult population of Germany. PLoS One. 2018;13(7):e0200267. doi:10.1371/journal.pone.0200267

31. UniProt, Consortium. UniProt: the universal protein knowledgebase in 2021. Nucleic Acids Res. 2021;49(D1):D480–D489. doi:10.1093/nar/gkaa1100

32. R Core Development Team. A Language and Environment for Statistical Computing,. R Foundation for Statistical Computing; 2019. https://www.R-project.org/

33. Michael L, Wolfgang H, Herv\’e P es, et al. Software for Computing and Annotating Genomic Ranges.; 2013. doi:10.1371/journal.pcbi.1003118

34. Dhanda SK, Vaughan K, Schulten V, et al. Development of a novel clustering tool for linear peptide sequences. Immunology. 2018;155(3):331–345. doi:10.1111/imm.12984

35. Wright E. DECIPHER: Tools for curating, analyzing, and manipulating biological sequences. Published online 2021.

36. Poh CM, Carissimo G, Wang B, et al. Two linear epitopes on the SARS-CoV-2 spike protein that elicit neutralising antibodies in COVID-19 patients. Nat Commun. 2020;11(1):2806. doi:10.1038/s41467-020-16638-2

37. Vanderheijden N, Stevaert A, Xie J, et al. Functional Analysis of Human and Feline Coronavirus Cross-Reactive Antibodies Directed Against the SARS-CoV-2 Fusion Peptide. Front Immunol. 2021;12:790415. doi:10.3389/fimmu.2021.790415

38. Sun X, Yi C, Zhu Y, et al. Neutralization mechanism of a human antibody with pan-coronavirus reactivity including SARS-CoV-2. Nat Microbiol. 2022;7(7):1063–1074. doi:10.1038/s41564-022-01155-3

39. Dacon C, Tucker C, Peng L, et al. Broadly neutralizing antibodies target the coronavirus fusion peptide. Science. 2022;377(6607):728-735. doi:10.1126/science.abq3773

40. Low JS, Jerak J, Tortorici MA, et al. ACE2-binding exposes the SARS-CoV-2 fusion peptide to broadly neutralizing coronavirus antibodies. Science. 2022;377(6607):735-742. doi:10.1126/science.abq2679

41. Le K, Kannappan S, Kim T, Lee JH, Lee HR, Kim KK. Structural understanding of SARS-CoV-2 virus entry to host cells. Front Mol Biosci. 2023;10:1288686. doi:10.3389/fmolb.2023.1288686

42. Xu GJ, Kula T, Xu Q, et al. Viral immunology. Comprehensive serological profiling of human populations using a synthetic human virome. Science. 2015;348(6239):aaa0698. doi:10.1126/science.aaa0698

43. Shrock E, Fujimura E, Kula T, et al. Viral epitope profiling of COVID-19 patients reveals cross-reactivity and correlates of severity. Science. 2020;370(6520). doi:10.1126/science.abd4250

44. Vita R, Mahajan S, Overton JA, et al. The Immune Epitope Database (IEDB): 2018 update. Nucleic Acids Res. 2019;47(D1):D339–D343. doi:10.1093/nar/gky1006

45. Amrun SN, Lee CY, Lee B, et al. Linear B-cell epitopes in the spike and nucleocapsid proteins as markers of SARS-CoV-2 exposure and disease severity. EBioMedicine. 2020;58:102911. doi:10.1016/j.ebiom.2020.102911

46. Simula ER, Manca MA, Jasemi S, et al. HCoV-NL63 and SARS-CoV-2 Share Recognized Epitopes by the Humoral Response in Sera of People Collected Pre- and during CoV-2 Pandemic. Microorganisms. 2020;8(12). doi:10.3390/microorganisms8121993

47. Lu S, Xie XX, Zhao L, et al. The immunodominant and neutralization linear epitopes for SARS-CoV-2. Cell Rep. 2021;34(4):108666. doi:10.1016/j.celrep.2020.108666

48. Kang S, Yang M, He S, et al. A SARS-CoV-2 antibody curbs viral nucleocapsid protein-induced complement hyperactivation. Nat Commun. 2021;12(1):2697. doi:10.1038/s41467-021-23036-9

49. Wang X, Lam JY, Chen L, et al. Mining of linear B cell epitopes of SARS-CoV-2 ORF8 protein from COVID-19 patients. Emerg Microbes Infect. 2021;10(1):1016–1023. doi:10.1080/22221751.2021.1931465

50. Prakash S, Srivastava R, Coulon PG, et al. Genome-Wide B Cell, CD4(+), and CD8(+) T Cell Epitopes That Are Highly Conserved between Human and Animal Coronaviruses, Identified from SARS-CoV-2 as Targets for Preemptive Pan-Coronavirus Vaccines. J Immunol. 2021;206(11):2566–2582. doi:10.4049/jimmunol.2001438

51. Zhang Y, Yang Z, Tian S, et al. A newly identified linear epitope on non-RBD region of SARS-CoV-2 spike protein improves the serological detection rate of COVID-19 patients. BMC Microbiol. 2021;21(1):194. doi:10.1186/s12866-021-02241-y

52. Pinto D, Sauer MM, Czudnochowski N, et al. Broad betacoronavirus neutralization by a stem helix-specific human antibody. Science. 2021;373(6559):1109-1116. doi:10.1126/science.abj3321

53. Patarroyo ME, Patarroyo MA, Alba MP, et al. The First Chemically-Synthesised, Highly Immunogenic Anti-SARS-CoV-2 Peptides in DNA Genotyped Aotus Monkeys for Human Use. Front Immunol. 2021;12:724060. doi:10.3389/fimmu.2021.724060

54. Polvere I, Voccola S, Parrella A, et al. A Peptide-Based Assay Discriminates Individual Antibody Response to the COVID-19 Pfizer/BioNTech mRNA Vaccine. Vaccines (Basel). 2021;9(9). doi:10.3390/vaccines9090987

55. Polyiam K, Phoolcharoen W, Butkhot N, et al. Immunodominant linear B cell epitopes in the spike and membrane proteins of SARS-CoV-2 identified by immunoinformatics prediction and immunoassay. Sci Rep. 2021;11(1):20383. doi:10.1038/s41598-021-99642-w

56. Gao F, Huang J, Li T, et al. A Highly Conserved Peptide Vaccine Candidate Activates Both Humoral and Cellular Immunity Against SARS-CoV-2 Variant Strains. Front Immunol. 2021;12:789905. doi:10.3389/fimmu.2021.789905

57. Li W, Chen Y, Prevost J, et al. Structural basis and mode of action for two broadly neutralizing antibodies against SARS-CoV-2 emerging variants of concern. Cell Rep. 2022;38(2):110210. doi:10.1016/j.celrep.2021.110210

58. Maghsood F, Shokri MR, Jeddi-Tehrani M, et al. Identification of immunodominant epitopes on nucleocapsid and spike proteins of the SARS-CoV-2 in Iranian COVID-19 patients. Pathog Dis. 2022;80(1). doi:10.1093/femspd/ftac001

59. Zhou P, Yuan M, Song G, et al. A human antibody reveals a conserved site on beta-coronavirus spike proteins and confers protection against SARS-CoV-2 infection. Sci Transl Med. 2022;14(637):eabi9215. doi:10.1126/scitranslmed.abi9215

60. Herrscher C, Eymieux S, Gaborit C, et al. ELISA-Based Analysis Reveals an Anti-SARS-CoV-2 Protein Immune Response Profile Associated with Disease Severity. J Clin Med. 2022;11(2). doi:10.3390/jcm11020405

61. Yu J, Qin Z, Liu X, et al. High-specificity targets in SARS-CoV-2 N protein for serological detection and distinction from SARS-CoV. Comput Biol Med. 2022;143:105272. doi:10.1016/j.compbiomed.2022.105272

62. Yang H, Cao J, Lin X, et al. Developing an Effective Peptide-Based Vaccine for COVID-19: Preliminary Studies in Mice Models. Viruses. 2022;14(3). doi:10.3390/v14030449

63. Tajuelo A, Lopez-Siles M, Mas V, et al. Cross-Recognition of SARS-CoV-2 B-Cell Epitopes with Other Betacoronavirus Nucleoproteins. Int J Mol Sci. 2022;23(6). doi:10.3390/ijms23062977

64. Geanes ES, LeMaster C, Fraley ER, et al. Cross-reactive antibodies elicited to conserved epitopes on SARS-CoV-2 spike protein after infection and vaccination. Sci Rep. 2022;12(1):6496. doi:10.1038/s41598-022-10230-y

65. Garanina E, Hamza S, Stott-Marshall RJ, et al. Antibody and T Cell Immune Responses to SARS-CoV-2 Peptides in COVID-19 Convalescent Patients. Front Microbiol. 2022;13:842232. doi:10.3389/fmicb.2022.842232

66. Pratesi F, Errante F, Pacini L, et al. A SARS-CoV-2 Spike Receptor Binding Motif Peptide Induces Anti-Spike Antibodies in Mice andIs Recognized by COVID-19 Patients. Front Immunol. 2022;13:879946. doi:10.3389/fimmu.2022.879946

67. Garrido JL, Medina MA, Bravo F, et al. IgG targeting distinct seasonal coronavirus-conserved SARS-CoV-2 spike subdomains correlates with differential COVID-19 disease outcomes. Cell Rep. 2022;39(9):110904. doi:10.1016/j.celrep.2022.110904

68. Yi Z, Ling Y, Zhang X, et al. Functional mapping of B-cell linear epitopes of SARS-CoV-2 in COVID-19 convalescent population. Emerg Microbes Infect. 2020;9(1):1988–1996. doi:10.1080/22221751.2020.1815591

69. Gregory DJ, Vannier A, Duey AH, et al. Repertoires of SARS-CoV-2 epitopes targeted by antibodies vary according to severity of COVID-19. Virulence. 2022;13(1):890–902. doi:10.1080/21505594.2022.2073025

70. Ng KW, Faulkner N, Cornish GH, et al. Preexisting and de novo humoral immunity to SARS-CoV-2 in humans. Science. 2020;370(6522):1339-1343. doi:10.1126/science.abe1107

71. Musico A, Frigerio R, Mussida A, et al. SARS-CoV-2 Epitope Mapping on Microarrays Highlights Strong Immune-Response to N Protein Region. Vaccines (Basel). 2021;9(1). doi:10.3390/vaccines9010035

72. Ladner JT, Henson SN, Boyle AS, et al. Epitope-resolved profiling of the SARS-CoV-2 antibody response identifies cross-reactivity with endemic human coronaviruses. Cell Rep Med. 2021;2(1):100189. doi:10.1016/j.xcrm.2020.100189

73. Holenya P, Lange PJ, Reimer U, et al. Peptide microarray-based analysis of antibody responses to SARS-CoV-2 identifies unique epitopes with potential for diagnostic test development. Eur J Immunol. 2021;51(7):1839–1849. doi:10.1002/eji.202049101

74. Voss C, Esmail S, Liu X, et al. Epitope-specific antibody responses differentiate COVID-19 outcomes and variants of concern. JCI Insight. 2021;6(13). doi:10.1172/jci.insight.148855

75. Mishra N, Huang X, Joshi S, et al. Immunoreactive peptide maps of SARS-CoV-2. Commun Biol. 2021;4(1):225. doi:10.1038/s42003-021-01743-9

76. Schwarz T, Heiss K, Mahendran Y, et al. SARS-CoV-2 Proteome-Wide Analysis Revealed Significant Epitope Signatures in COVID-19 Patients. Front Immunol. 2021;12:629185. doi:10.3389/fimmu.2021.629185

77. Heffron AS, McIlwain SJ, Amjadi MF, et al. The landscape of antibody binding in SARS-CoV-2 infection. PLoS Biol. 2021;19(6):e3001265. doi:10.1371/journal.pbio.3001265

78. Nitahara Y, Nakagama Y, Kaku N, et al. High-Resolution Linear Epitope Mapping of the Receptor Binding Domain of SARS-CoV-2 Spike Protein in COVID-19 mRNA Vaccine Recipients. Microbiol Spectr. 2021;9(3):e0096521. doi:10.1128/Spectrum.00965-21

79. Camerini D, Randall AZ, Trappl-Kimmons K, et al. Mapping SARS-CoV-2 Antibody Epitopes in COVID-19 Patients with a Multi-Coronavirus Protein Microarray. Microbiol Spectr. 2021;9(2):e0141621. doi:10.1128/Spectrum.01416-21

80. Gattinger P, Niespodziana K, Stiasny K, et al. Neutralization of SARS-CoV-2 requires antibodies against conformational receptor-binding domain epitopes. Allergy. 2022;77(1):230–242. doi:10.1111/all.15066

81. Li M, Liu J, Lu R, et al. Longitudinal immune profiling reveals dominant epitopes mediating long-term humoral immunity in COVID-19-convalescent individuals. J Allergy Clin Immunol. 2022;149(4):1225–1241. doi:10.1016/j.jaci.2022.01.005

82. Levy Y, Alcalay R, Zvi A, et al. Immunodominant Linear B-Cell Epitopes of SARS-CoV-2 Spike, Identified by Sera from K18-hACE2 Mice Infected with the WT or Variant Viruses. Vaccines (Basel). 2022;10(2). doi:10.3390/vaccines10020251

83. Chen L, Pang P, Qi H, et al. Evaluation of Spike Protein Epitopes by Assessing the Dynamics of Humoral Immune Responses in Moderate COVID-19. Front Immunol. 2022;13:770982. doi:10.3389/fimmu.2022.770982

84. Hotop SK, Reimering S, Shekhar A, et al. Peptide microarrays coupled to machine learning reveal individual epitopes from human antibody responses with neutralizing capabilities against SARS-CoV-2. Emerg Microbes Infect. 2022;11(1):1037–1048. doi:10.1080/22221751.2022.2057874

85. Stoddard CI, Galloway J, Chu HY, et al. Epitope profiling reveals binding signatures of SARS-CoV-2 immune response in natural infection and cross-reactivity with endemic human CoVs. Cell Rep. 2021;35(8):109164. doi:10.1016/j.celrep.2021.109164

86. Guo JY, Liu IJ, Lin HT, et al. Identification of COVID-19 B-cell epitopes with phage-displayed peptide library. J Biomed Sci. 2021;28(1):43. doi:10.1186/s12929-021-00740-8

87. Tang J, Lee Y, Ravichandran S, et al. Epitope diversity of SARS-CoV-2 hyperimmune intravenous human immunoglobulins and neutralization of variants of concern. iScience. 2021;24(9):103006. doi:10.1016/j.isci.2021.103006

88. Li L, Gao M, Li J, et al. Methods to Identify Immunogenic Peptides in SARS-CoV-2 Spike and Protective Monoclonal Antibodies in COVID-19 Patients. Small Methods. 2021;5(7):2100058. doi:10.1002/smtd.202100058

89. Ravichandran S, Tang J, Grubbs G, et al. SARS-CoV-2 immune repertoire in MIS-C and pediatric COVID-19. Nat Immunol. 2021;22(11):1452–1464. doi:10.1038/s41590-021-01051-8

90. Garrett ME, Galloway JG, Wolf C, et al. Comprehensive characterization of the antibody responses to SARS-CoV-2 Spike protein finds additional vaccine-induced epitopes beyond those for mild infection. Elife. 2022;11. doi:10.7554/eLife.73490

91. Kumar G, Sterrett S, Hall L, et al. Comprehensive mapping of SARS-CoV-2 peptide epitopes for development of a highly sensitive serological test for total and neutralizing antibodies. Protein Eng Des Sel. 2022;35. doi:10.1093/protein/gzab033

92. Willcox AC, Sung K, Garrett ME, et al. Detailed analysis of antibody responses to SARS-CoV-2 vaccination and infection in macaques. PLoS Pathog. 2022;18(4):e1010155. doi:10.1371/journal.ppat.1010155

93. Ballmann R, Hotop SK, Bertoglio F, et al. ORFeome Phage Display Reveals a Major Immunogenic Epitope on the S2 Subdomain of SARS-CoV-2 Spike Protein. Viruses. 2022;14(6). doi:10.3390/v14061326

94. Haynes WA, Kamath K, Bozekowski J, et al. High-resolution epitope mapping and characterization of SARS-CoV-2 antibodies in large cohorts of subjects with COVID-19. Commun Biol. 2021;4(1):1317. doi:10.1038/s42003-021-02835-2

95. Zamecnik CR, Rajan JV, Yamauchi KA, et al. ReScan, a Multiplex Diagnostic Pipeline, Pans Human Sera for SARS-CoV-2 Antigens. Cell Rep Med. 2020;1(7):100123. doi:10.1016/j.xcrm.2020.100123

96. Wang H, Wu X, Zhang X, et al. SARS-CoV-2 Proteome Microarray for Mapping COVID-19 Antibody Interactions at Amino Acid Resolution. ACS Cent Sci. 2020;6(12):2238–2249. doi:10.1021/acscentsci.0c00742

97. Jones BE, Brown-Augsburger PL, Corbett KS, et al. The neutralizing antibody, LY-CoV555, protects against SARS-CoV-2 infection in nonhuman primates. Sci Transl Med. 2021;13(593). doi:10.1126/scitranslmed.abf1906

98. Sen SR, Sanders EC, Gabriel KN, et al. Predicting COVID-19 Severity with a Specific Nucleocapsid Antibody plus Disease Risk Factor Score. mSphere. 2021;6(2). doi:10.1128/mSphere.00203-21

99. Woudenberg T, Pelleau S, Anna F, et al. Humoral immunity to SARS-CoV-2 and seasonal coronaviruses in children and adults in north-eastern France. EBioMedicine. 2021;70:103495. doi:10.1016/j.ebiom.2021.103495

100. Lv H, Wu NC, Tsang OT, et al. Cross-reactive Antibody Response between SARS-CoV-2 and SARS-CoV Infections. Cell Rep. 2020;31(9):107725. doi:10.1016/j.celrep.2020.107725

101. Song G, He WT, Callaghan S, et al. Cross-reactive serum and memory B-cell responses to spike protein in SARS-CoV-2 and endemic coronavirus infection. Nat Commun. 2021;12(1):2938. doi:10.1038/s41467-021-23074-3

102. Dacon C, Peng L, Lin TH, et al. Rare, convergent antibodies targeting the stem helix broadly neutralize diverse betacoronaviruses. Cell Host Microbe. 2022;31(1):97–111.e12. doi:10.1016/j.chom.2022.10.010

103. Jia L, Weng S, Wu J, et al. Preexisting antibodies targeting SARS-CoV-2 S2 cross-react with commensal gut bacteria and impact COVID-19 vaccine induced immunity. Gut Microbes. 2022;14(1):2117503. doi:10.1080/19490976.2022.2117503

104. Castleman MJ, Stumpf MM, Therrien NR, et al. SARS-CoV-2 infection relaxes peripheral B cell tolerance. J Exp Med. 2022;219(6). doi:10.1084/jem.20212553

105. Jiang W, Johnson D, Adekunle R, et al. COVID-19 is associated with bystander polyclonal autoreactive B cell activation as reflected by a broad autoantibody production, but none is linked to disease severity. J Med Virol. 2023;95(1):e28134. doi:10.1002/jmv.28134

106. Center for Systems Science and Engineering (CSSE). COVID-19 Data Repository by the Center for Systems Science and Engineering (CSSE) at Johns Hopkins University. Published online March 26, 2020. Accessed February 20, 2024. https://github.com/CSSEGISandData/COVID-19/blob/master/archived_data/archived_time_series/time_series_19-covid-Confirmed_archived_0325.csv

107. Bach E, Fitzgerald SF, Williams-MacDonald SE, et al. Genome-wide epitope mapping across multiple host species reveals significant diversity in antibody responses to Coxiella burnetii vaccination and infection. Front Immunol. 2023;14:1257722. doi:10.3389/fimmu.2023.1257722

108. Lo KC, Sullivan E, Bannen RM, et al. Comprehensive Profiling of the Rheumatoid Arthritis Antibody Repertoire. Arthritis Rheumatol. 2020;72(2):242–250. doi:10.1002/art.41089

